# Subpopulation-specific gene expression in *Lachancea thermotolerans* uncovers distinct metabolic adaptations to wine fermentation

**DOI:** 10.1101/2024.09.05.611386

**Authors:** Javier Vicente, Santiago Benito, Domingo Marquina, Antonio Santos

## Abstract

Gene expression is the first step in translating genetic information into quantifiable traits. This study analysed gene expression in 23 strains across six subpopulations of *Lachancea thermotolerans*, shaped by anthropization, under winemaking conditions to understand the impact of adaptation on transcriptomic profiles and fermentative performance, particularly regarding lactic acid production. By sequencing mRNA during exponential growth and fermentation in synthetic grape must, we identified unique expression patterns linked to the strains originated from wine-related environments. Global expression analysis revealed that anthropized subpopulations, particularly Europe/Domestic-2 and Europe-Mix, exhibited distinct gene expression profiles related to fermentation processes such as glycolysis and pyruvate metabolism. These processes were differentially expressed, along with other important biological processes during fermentation, such as nitrogen and fatty acid metabolism. This study highlights that anthropization has driven metabolic specialization in *L. thermotolerans*, enhancing traits like lactic acid production, which is a trait of interest in modern winemaking. Correlation analysis further linked lactic acid dehydrogenase genes with key metabolic pathways, indicating adaptive gene expression regulation. Additionally, differences in other metabolites of oenological interest as glycerol or aroma compounds production are highlighted. Here, we provide insights into the evolutionary processes shaping the transcriptomic diversity of *L. thermotolerans*, emphasizing the impact of winemaking environments on driving specific metabolic adaptations, including lactic acid production. Understanding the gene expression differences linked to lactic acid production could allow a more rational address of biological acidification while optimizing yeast-specific nutritional requirements during fermentation.

**GRAPHICAL ABSTRACT:** 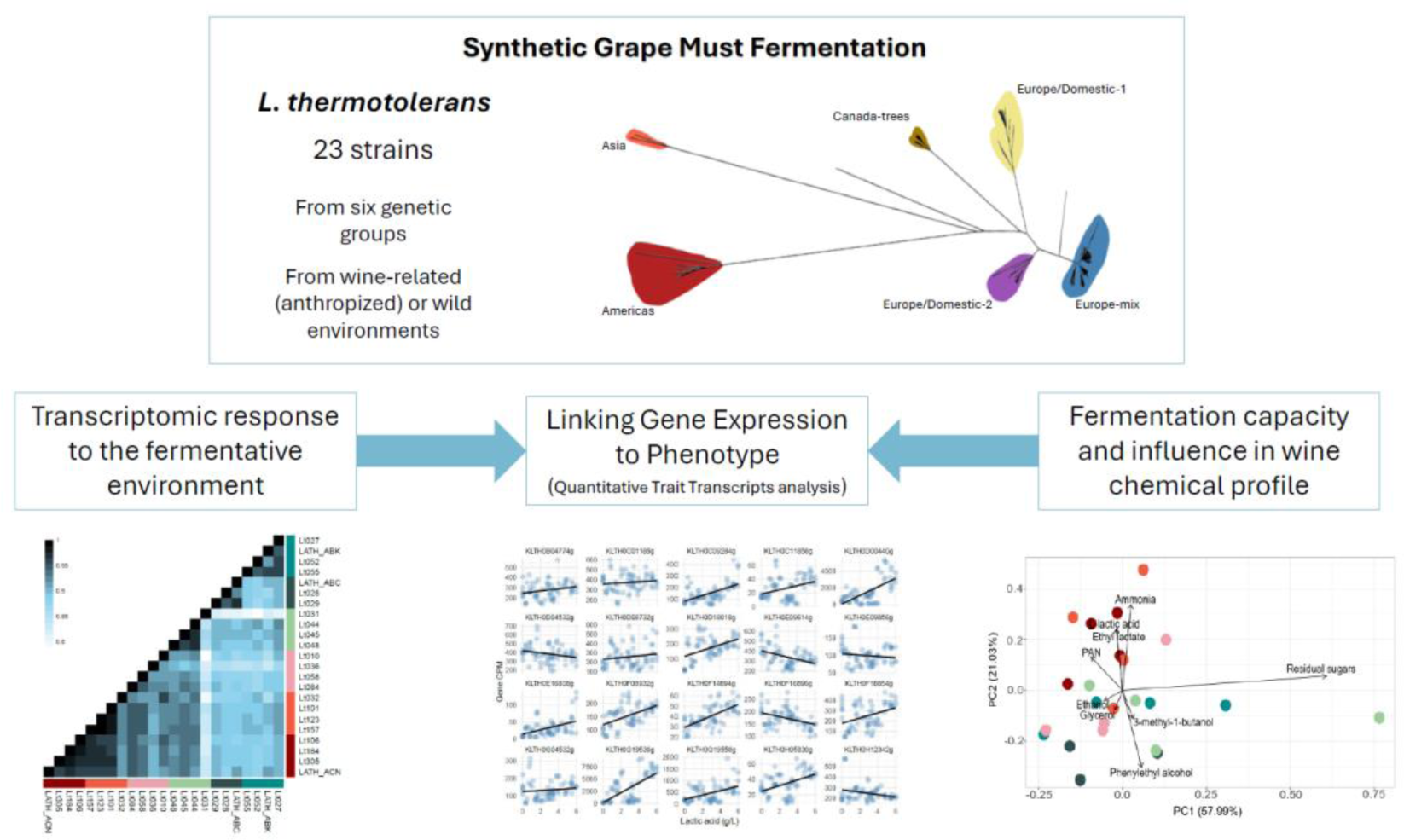

## 1. INTRODUCTION

Gene expression is the first step in which genetic information is transformed into a phenotype. Phenotypic variation within a certain species is the consequence of both genetic variants and expression regulation (Caudal et al., 2024). Divergence at the transcriptomic level represents one of the main drivers of the observed phenotypic diversity within a species, together with the genetic variants that originate it and the evolutionary processes that have shaped it (Jallet et al., 2023). It is often challenging to link observed genetic variation (in sequence or structure) to a specific phenotype (Villarreal et al., 2024). Expression analysis studies bridge this gap, as the phenotypic trait is a direct consequence of the expression profile. Generally, gene expression among different individuals from various genetic backgrounds and evolutionary origins evolves neutrally, showing few changes in gene expression profiles. However, studying the differential regulation of gene expression between individuals or genetic backgrounds can help in understanding the adaptive response as a consequence of the evolutionary forces that have driven the species evolution (Jallet et al., 2023).

Grape must is a challenging medium for microorganism development due to several factors, including high osmotic pressures, nutrient scarcity (such as nitrogen, vitamins, or minerals as cofactors), the presence of antimicrobials (SO₂, Cu or chitosan), and acidic pH. The competition for nutrients, the physicochemical conditions and various biotic factors, create an evolutionary pressure that has greatly affected the inhabiting microorganisms, particularly wine-related yeasts. Yeasts have evolved in response to this specific environment, with certain metabolic traits emerging to make them more competitive. The basis of all these phenotypic traits results from, among others, the increase in gene copy numbers, chromosomal rearrangements, the acquisition of new genes through horizontal gene transfer, or the regulation of expression levels (Becerra-Rodríguez et al., 2020; García-Ríos & Guillamón, 2022; Jallet et al., 2023; Marsit & Dequin, 2015; Villarreal et al., 2022). Additionally, the anaerobic capability and Crabtree effect emerged around 150 and 100 million years ago, respectively, in response to high sugars environments. This new metabolic trait allowed for enhanced sugar consumption and antimicrobial production (i.e. ethanol), providing greater fitness in those environments (Hagman et al., 2013).

Initially, these were fermentative-wild environments, but they evolved alongside human culture, transforming into industrial and human-controlled ones. Winemaking dates back to 6,500 years ago and has shaped the evolution of several species related to this environment, including not only *Saccharomyces cerevisiae* but also other yeasts such as *Torulaspora delbrueckii, Hanseniaspora uvarum, Brettanomyces bruxellensis*, and *Lachancea thermotolerans* (Albertin et al., 2016; Gounot et al., 2020; Ruiz et al., 2021; Silva et al., 2023). Interest in *L. thermotolerans* has grown recently, with research indicating the influence of anthropization on its evolution. Previous insights into this yeast’s diversity have been published over the last decade (Freel et al., 2014; Hranilovic et al., 2017; Vicente, Friedrich, et al., 2024). Today, interest in *L. thermotolerans* is driven primarily by its use as a biological approach for acidity management during wine fermentation. Climate change has introduced several challenges to modern oenological industry, including high ethanol concentrations, high pH, and low levels of malic acid. These issues have prompted researchers to explore alternative methods to mitigate potential risks and improve fermentation efficiency (Vicente et al., 2022). *L. thermotolerans*, through its ability to produce significant amounts of lactic acid from pyruvate concurrently with alcoholic fermentation, emerges as a promising alternative. This trait has been demonstrated to be not only strain-dependent but also linked to specific genomic clusters that have emerged due to anthropization. This trait, considered an anthropization signature, is seen as an adaptive step to support and adapt to the wine niche and fermentative environment by alleviating excess glycolytic flux (Vicente, Friedrich, et al., 2024). Thus, anthropized strains are believed to have modified their metabolic regulation as a result of adaptation to the fermentative environment through this alternative metabolic pathway allowing a faster regeneration of cofactors (Monnin et al., 2024; Shekhawat et al., 2020; Tyibilika et al., 2024). Furthermore, it has been previously shown that different strains exhibit different transcriptional responses to winemaking conditions (Battjes et al., 2023).

Considering that strains of *L. thermotolerans* exhibit different behaviours during their growth under winemaking conditions and produce significantly different amounts of lactic acid, these variations cannot be fully explained by the presence of distinct sets of genes encoding lactate dehydrogenases in the different strains. Instead, they suggest differences in overall metabolic regulation arising from differences in gene expression.

In this work, we have employed a representative collection of isolates from different genetic backgrounds representing the various subpopulations defining the species: Asia, Americas, Canada-trees, Europe/Domestic-1, Europe/Domestic-2 and Europe-Mix (Hranilovic et al., 2017; Vicente, Friedrich, et al., 2024). These groups show significant phenotypic variations, especially concerning lactic acid production. Three non-European subpopulations consist essentially of wild isolates, whereas the other three, originating mainly from Europe and linked to the Mediterranean basin—a traditional winemaking region—come from anthropized environments associated with wine-related environments. The selected strains enable us to determine how strains from different environments cope with the fermentative environment, revealing different transcriptional signatures and specific responses as a result of the adaptation process.

Our understanding of the connection between the response to challenging fermentative environments and genetic backgrounds in yeasts remains limited, although numerous examples have been reported (Caudal et al., 2024; Jallet et al., 2023; Lairón-Peris et al., 2020; Villarreal et al., 2024). This gap in knowledge highlights the complexity of gene expression regulation in yeast and the need for more comprehensive studies to determine how genetic variations influence both fermentation efficiency and product quality. Understanding these interactions is crucial for improving the production of wine and other fermented products, enabling the development of more robust and high-performing yeast strains suited to specific industrial needs or the specific requirements of each strain and species during the fermentation.

## 2. MATERIALS AND METHODS

### 2.1. Strains, culture media, cultures and sampling

For this study, we used 23 strains of *L. thermotolerans* selected from a previously characterised collection (Vicente, Friedrich, et al., 2024) (Table S1). Strains were selected based on the mean lactic acid production of each genomic subpopulation previously defined; the four strains closest to the mean production of each subpopulation (three in the case of the Asian subpopulation) were chosen as representatives of the genetic group. This approach ensured differential lactic acid production across the entire dataset and characteristic lactic acid production within each subpopulation. All strains were cryopreserved at -80 °C in 25 % glycerol and maintained on Sabouraud agar at 28 °C for daily handling.

Fermentation assays were performed in triplicate using 100 mL borosilicate bottles containing 95 mL of Synthetic Grape Must (SGM) (180 g/L of equimolecular glucose and fructose, 140 mg N/L from amino acids, 60 mg N/L from di-ammonium phosphate, pH 3.3), prepared as previously described (Henschke & Jiranek, 1993), and maintained at 25 °C and 100 rpm. Strains were inoculated, at their exponential growth phase, into the laboratory-scale fermentations at a final OD_600nm_ of 0.2 (10^6^ CFU/mL) from a 20 mL SGM preculture in 50 mL borosilicate bottles (28 °C,150 rpm, 16 hours). Fermentation kinetics were monitored by measuring weight loss (to three significant decimals), with fermentation considered complete when weight loss remained at or below 0.01 % over 24 hours for two consecutive days.

Samples (1.0 mL) were taken at different time points for different analyses. For CFU determination, samples were serially diluted 10-fold and plated on Sabouraud agar. For metabolite analysis, samples were centrifuged at 7,500 rpm for 5 minutes, and the supernatant was stored at -20°C until further analysis. For RNAseq, 1.7 mL samples were collected in quadruplicate at 30 hours of culture, flash-frozen in liquid nitrogen, and stored at −80°C until RNA extraction.

### 2.2. mRNA extractions, library preparation and sequencing

Total RNA was extracted from one of the samples stored at -80 °C using the Quick-RNA Fungal/Bacterial MicroPrep kit (Zymo Research, USA). The quality of the RNA was evaluated with a Bioanalyzer 2100 (Agilent Technologies, USA), and samples with an RNA Integrity Number (RIN) ≥ 7.5 were selected for further analysis. Directional mRNA libraries were prepared using polyA enrichment, and their quality was validated with a Qubit 3.0 (Thermo Fisher, USA) and a 2100 Bioanalyzer (Agilent Technologies, USA). The libraries were then sequenced on an Illumina NovaSeq X Plus sequencer using 150 bp paired-end sequencing, with an output of 3 Gb per sample.

### 2.3. RNAseq data processing and analysis of differentially expressed genes (DEGs)

Illumina paired-end reads were quality assessed with FastQC and trimmed using *Trimmomatic* (v.0.39) (Bolger et al., 2014). HISAT2 pipeline (v.2.2.1) (Kim et al., 2019) was employed to align the reads to the indexed reference genome *L. thermotolerans* CBS 6340 (Souciet et al., 2009) and to assemble the alignments into full-length transcripts. Counting of reads mapping to each genomic feature was performed using *featureCounts* (v.2.0.6) (Liao et al., 2014).

The differential expression analysis was performed on read counts using the *DESeq2* R package (v.1.44.0) (Love et al., 2014). Library size normalization was performed with the default method implemented by DESeq2. Expression in each subpopulation was contrasted against expression in samples from all other subpopulations considered as a whole. Each gene was considered differentially expressed if their false discovery rate (FDR)-corrected P-values were below 0.05 and the absolute log2 Fold-Change (FC) above 1 (corresponding to an upregulation or downregulation by a minimum of 2-fold)

### 2.4. Functional enrichment analysis

To conduct functional annotation of the DEGs, we performed Kyoto Encyclopedia of Genes and Genomes (KEGG) enrichment analyses using the *enrichKEGG* function from the *clusterProfiler* R package (v.4.12.2) (Wu et al., 2021). Enrichment terms were considered significant if their corrected P-adjusted value was below 0.05.

### 2.5. Identification of Cluster Specific transcriptomic Signatures

Cluster-Specific transcriptomic Signatures (CSS) were analysed as previously described (Jallet et al., 2023). These signatures represent transcriptomic profiles significantly over- or under-expressed within specific subpopulations. Initially, all DEGs with an absolute log2 fold-change above 1.6 (equivalent to a 3-fold change in expression) and an adjusted P-value ranking within the top half of differentially expressed genes (either upregulated or downregulated) were retained. Subsequently, CSS candidates were further refined by filtering out those present in more than one subpopulation.

### 2.6. Analysis of genes specifically linked to glycolytic and fermentative pathways

To focus on genes associated with glucose and lactic acid metabolism, pathway annotations from the SGD database were utilized (https://pathway.yeastgenome.org/, last accessed August 2024). *S. cerevisiae* genes belonging to “Glycolysis I (from glucose 6-phosphate)”, “Superpathway of glucose fermentation”, “TCA cycle, aerobic respiration”, and “aerobic respiration, electron transport chain” subsets were retrieved and queried against *L. thermotolerans* to identify orthologs. Additionally, all genes from the *L. thermotolerans* glycolysis/gluconeogenesis KEGG pathway were included (https://www.genome.jp/kegg/, last accessed August 2024). For specific genes correlations, Spearman’s rank correlation and associated p-values were computed using the *Hmisc* R package (v.5.1-3) (Harrell Jr & Harrell Jr, 2019) for each gene pair. Genes networks were constructed using the significant correlations (p < 0.05) in *igraph* R package (v. 2.0.3) (Csárdi et al., 2023).

### 2.7. Metabolite profiling of the fermentations

The concentrations of different metabolites were quantified at different fermentation stages. Total sugars (glucose and fructose), glycerol, L-lactic acid, L-malic acid, acetic acid, ammonia, and Primary Amino Nitrogen (PAN) were enzymatically quantified using a Y15-autoanalyzer (Biosystems, Spain). Additionally, at the end of the fermentation, pH and total acidity were determined by FTIR analysis using a Bacchus 3 MultiSpec (Tecnología de Difusión Ibérica, Spain). An analysis of the volatile fraction of the resulting wines was conducted according to previously described methodology using liquid-liquid microextraction followed by GC-MS (Coelho et al., 2020). All statistical analysis were carried out using R. Spearman’s correlation and associated p-values were computed using the Hmisc R package (v.5.1-3) (Harrell Jr & Harrell Jr, 2019) for each metabolite measured at the end of the fermentation.

### 2.8. Quantitative trait transcripts linked to lactic acid production

We used a linear regression model to identify Quantitative Trait Transcripts (QTTs) correlating with lactic acid production, based on previously described methods (Kong et al., 2022; Pang et al., 2019). Hidden factors were extracted by a Surrogate Variable Analysis (SVA) method, using *sva* R package (v.3.52.0) (Leek JT, 2024) with a model matrix created based on the metabolomic data, which were included as predictors in the regression models. For each gene, a linear regression model was fitted to examine the relationship between gene expression and lactic acid values. The model used was:

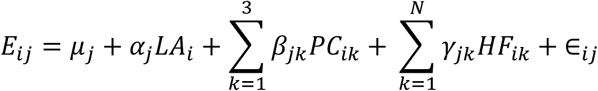

where *E*_*ij*_ represents the expression level of gene *j* in sample *i*, μ_*j*_ is the regression intercept, *LA*_*i*_ denotes the lactic acid level in sample *i*, *PC*_*ik*_ (1 ≤ *k* ≤ 3) indicates the value of the *k*-th principal component for sample *i*, *HF*_*ik*_ (1 ≤ *k* ≤ *N*) indicates the value of the *k*-th hidden factor for sample *i*, *N* is the number of hidden factors considered, ε_*ij*_ is the error term, and α_*j*_, β_*jk*_ and γ_*jk*_ are the regression coefficients for lactic acid levels, the *k*-th principal component, and the *k*-th hidden factor, respectively. For multiple testing, the p-values were adjusted using the Benjamini-Hochberg method using *stats* R package. Genes with adjusted p-values less than or equal to 0.001 were defined as QTTs. Among these, genes with positive regression coefficients (α_*j*_>0) were classified as positively correlated with lactic acid production, while those with negative coefficients (α_*j*_<0) were classified as negatively correlated.

## 3. RESULTS

To determine the expression patterns within *L. thermotolerans*, we selected a set of 23 strains representing the following subpopulations: Americas, Asia, Canada-trees, Europe/Domestic-1, Europe/Domestic-2, and Europe-mix. These subpopulations are characterized both by geographical origin and ecological niche, with Asia, Americas and Canada-trees comprising strains from wild environments, and Europe/Domestic-1, Europe/Domestic-2, and Europe-mix consisting of anthropized strains isolated from vineyards and grape must (Hranilovic et al., 2017; Vicente, Friedrich, et al., 2024). Our previous research highlighted the significant impact of anthropization on species evolution, influencing genetic differentiation, as revealed by whole-genome sequencing, and phenotypic traits such as antimicrobial resistance, carbon/nitrogen source utilization, and particularly, lactic acid production during fermentation (Vicente, Friedrich, et al., 2024). In this study, we aimed to elucidate the genetic regulation of these phenotypic variations by analyzing transcriptomic profiles under winemaking conditions. We selected strains based on their ability to produce lactic acid, targeting those closest to the subpopulation previously defined mean levels. RNA was extracted at 30 hours of fermentation, a time when fermentation kinetics are in exponential phase, maximum cell density has been reached, and lactic acid production shows the highest rates (Battjes et al., 2023). Additionally, we conducted metabolome analyses to determine its impact in the fermentation process and its metabolic outcomes at several timepoints of the fermentation.

### 3.1. Global view of gene expression

To assess gene expression variation in *L. thermotolerans*, we sequenced the total mRNA from a previously characterized set of 23 isolates during exponential growth and fermentation kinetics in a defined synthetic grape must. *L. thermotolerans* CBS 6340 was used as the reference for direct mapping of reads. Based on the aligned reads, Principal Component Analysis (Figure 1a) of Counts Per Million (CPM) for each gene and strain revealed two distinct groups of samples corresponding to the previously defined wild and anthropized subpopulations (Table S2). Within the anthropized subpopulations, strains from Europe/Domestic-2 grouped together with Europe-Mix, while Europe/Domestic-1, Canada-trees, Americas, and Asia subpopulations group together, exhibiting more distinct expression patterns under wine fermentation conditions. This grouping strongly suggests that the anthropization process has influenced the adaptation and gene expression patterns of this species in the winemaking environment.

**Figure 1.**
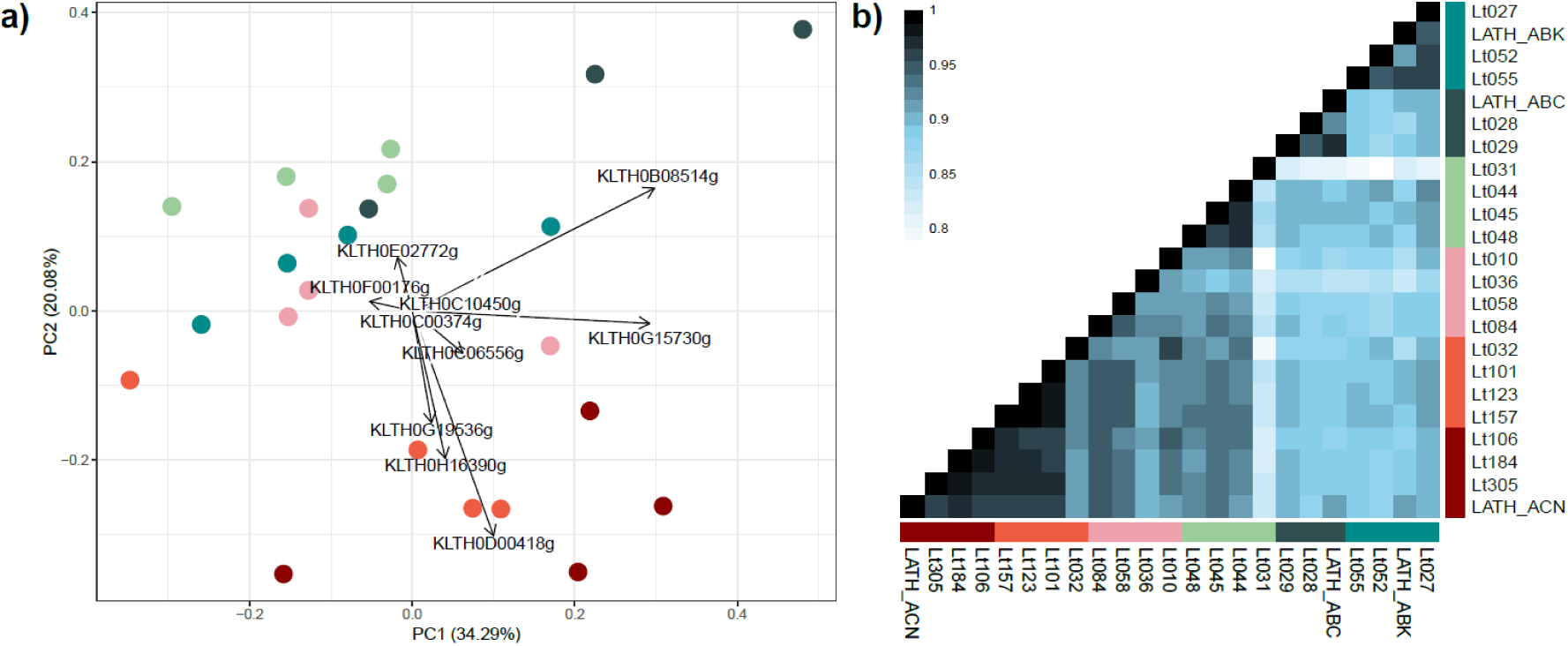
Distinctive patterns of global expression profiles across populations. a) Principal Component Analysis (PCA) of the mean copies per million (CPM) of each gene and strain. b) Pairwise correlation matrix between isolates expression profiles. Correlation coefficient was computed using the Spearman correlation. Samples were intendedly ordered by population. Different colours represent the different subpopulations: Asia: dark green; Americas: blue; Canada-trees: light green; Europe/Domestic-1: pink; Europe/Domestic-2: orange; Europe-mix: brown.

Notable differences were observed across isolates from different environments. Regarding specific genes, among the top ten main drivers of the transcriptomic profiles’ distribution, we identified key enzymes involved in alcoholic fermentation, including glycolysis. Specifically, PDC (KLTH0D00418) and LDH (KLTH0G19536) showed higher transcript levels in Europe/Domestic-2 and Europe-Mix, whereas *HXT6* (KLTH0E02772) exhibited increased expression in the other subpopulations. Other genes contributing to the intra-group distribution include *ENO1* (KLTH0B08514) and TDH (KLTH0G15730), which encode enolase and glyceraldehyde-3-phosphate dehydrogenase activities, respectively—both essential enzymes in glycolysis.

To obtain a better understanding of the relationships between different transcriptomes, we performed a correlational analysis of the expression profiles across samples. In the pairwise correlation matrix based on whole-genome expression profiles, significant relationships were observed (Figure 1b). Samples from the Americas and Asian subpopulations exhibited the lowest correlation indices when compared to the rest of the samples, whereas the Canada-trees subpopulation, also considered a wild group, showed high correlation indices with the anthropized strains’ expression profiles. Notably, the highest correlation indices were observed between strains from the Europe/Domestic-2 and Europe-Mix subpopulations, further confirming the role of anthropization and niche adaptation in shaping the expression profiles of these strains. Interestingly, Lt031, a strain isolated from grapes (and considered anthropized) within the Canada-trees subpopulation, exhibited both the lowest intra- and inter-subpopulation correlation indices in the entire dataset.

### 3.2. Subpopulation-level expression variation: analysis of DEGs and functional enrichment

The Differential Expression (DE) analysis comparing each subpopulation to the rest of the samples identified an average of 225 DEGs across the six populations, with values ranging from 79 (Europe/Domestic-2) to 388 (Asia and Americas), according to the specified thresholds (absolute FC > 1 and adjusted p-value < 0.05) (Table 1). The direction of these DEGs varied between clusters. The general trend indicated downregulation (negative FC values), with a mean of approximately 161 downregulated genes, ranging from 28 in Europe/Domestic-1 to 321 in the Americas subpopulation. Upregulated genes were less common, with a mean of 64 upregulated genes, ranging from 26 in Europe/Domestic-2 to 105 in Europe/Domestic-1. Notably, significant FC values were observed among the downregulated genes in the wild subpopulations, with several genes showing FC values between 20 and 30, and mean FC values around -3.30. In contrast, anthropized clusters exhibited mean FC values around -1.90 for downregulated genes.

**Table 1.**
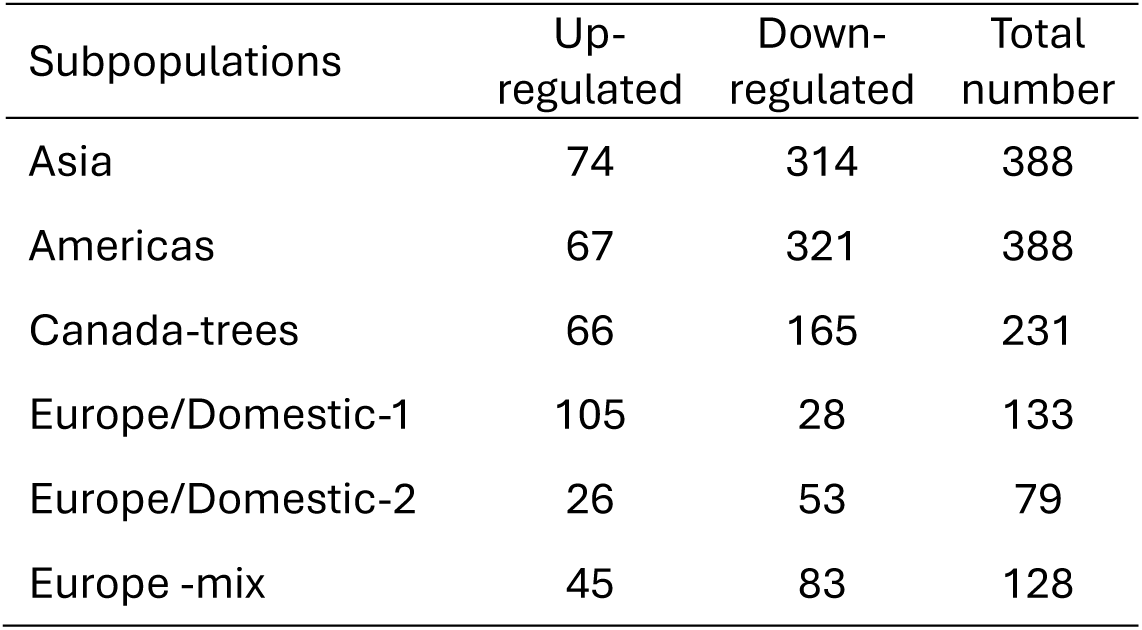
Differentially Expressed Genes (DEGs) by the different subpopulations. Only significative genes (p<0.05) with FC >1 were considered.

The KEGG pathway enrichment unveiled the biological processes that distinctly set the populations from each other (Figure 2, Table S3). DEGs were notably enriched in those associated with glycolysis and pyruvate metabolism, nitrogen metabolism, and fatty acid degradation and biosynthesis. The analysis of the accumulated FC for each biological process revealed significant differences in carbon metabolism (glycolysis and pyruvate metabolism) between subpopulations. Genes involved in these processes were generally downregulated in the Asia, Canada-trees, and Europe/Domestic-1 strains, while the anthropized clusters Europe/Domestic-2 and Europe-Mix exhibited notable upregulation of these genes. Among them, we found genes related with alcoholic fermentation as *PDC5* (KLTH0D00418), *PDC1* (KLTH0D08272) or the three LDH genes in *L. thermotolerans* (KLTH0D00440, KLTH0G19536, KLTH0G19558).

**Figure 2.**
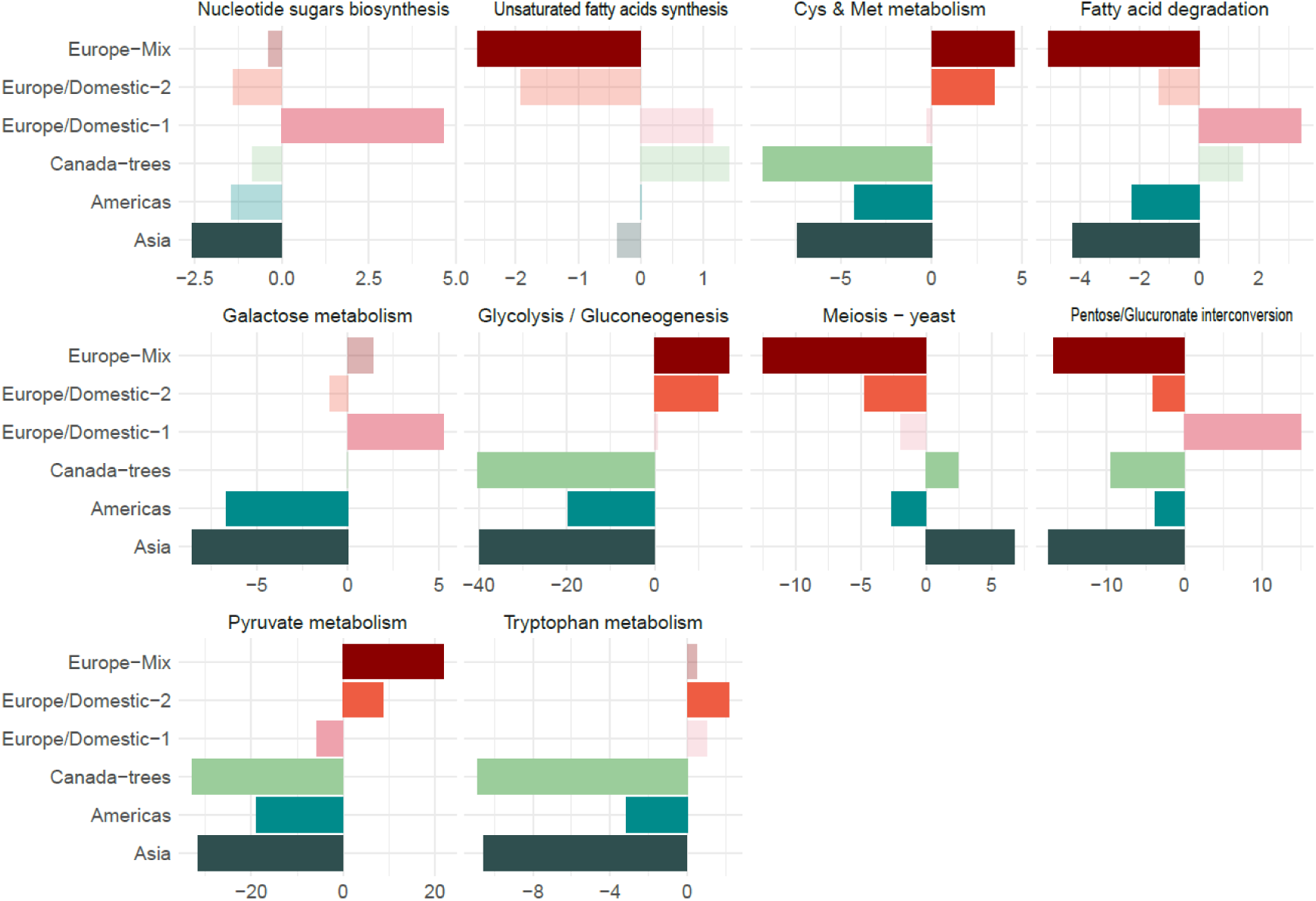
KEGG enrichment analysis for DEGs in each population, represented as the sum of accumulated fold changes (FC) for each category. All enriched terms found in any of the subpopulations are shown (in bold if found in the specific subpopulation, in light if the term was not specifically enriched). The colour code for the subpopulations is consistent with that used in Figure 1.

The overall accumulated FC across the subpopulations revealed a diverse landscape. Asia and Canada-trees exhibited accumulated downregulated values near -40 for glycolysis (-30 for pyruvate metabolism) enriched genes, compared to a positive FC near 15 in Europe/Domestic-2 and Europe-mix. Regarding nitrogen metabolism, significant differences were observed in cysteine, methionine, and tryptophan metabolism. These processes were primarily downregulated in the Canada-trees subpopulation. However, relevant differences emerged, particularly in the cysteine and methionine metabolism gene sets when calculating the accumulated FC across all subpopulations. The Americas, Asia, and Canada-trees subpopulations exhibited accumulated FC values near -5, while Europe/Domestic-2 and Europe-mix displayed accumulated FC values around 4, indicating distinct strategies in carbon and nitrogen metabolism among the different defined subpopulations.

### 3.3. Subpopulations specific response to the fermentative condition

To characterize CSS of each subpopulation, we identified genes displaying strong over- or under-expression in each subpopulation compared to the others. We selected genes showing a 3-fold variation with a low p-adjusted value (within the bottom half of the whole dataset). These criteria allowed us to identify a varying number of unique CSS for each subpopulation: Asia, 40; Americas, 43; Canada-trees, 11; Europe/Domestic-1, 5; Europe/Domestic-2, 2; Europe-Mix, 3 (Figure 3a, Table S4).

**Figure 3.**
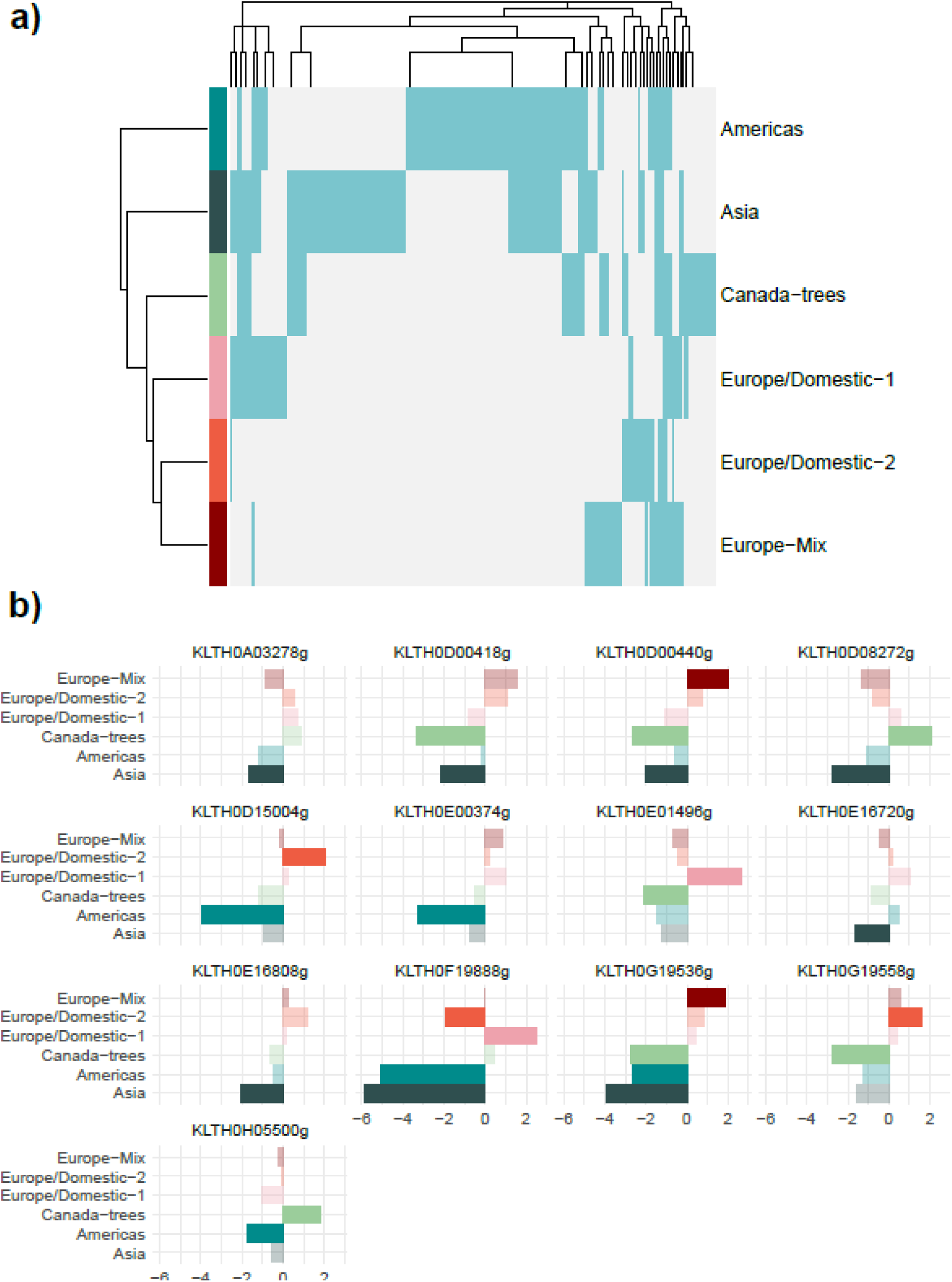
Cluster-Specific Signatures (CSS) differentiating subpopulations of *L. thermotolerans.* a) Presence matrix of all CSS candidates across the different strains. b) Fold change of selected CSS in the different subpopulations (in bold if found in the specific subpopulation, in light if the gene was not specifically enriched). The colour code for the subpopulations is consistent with that used in Figure 1.

When evaluating all CSS candidates (genes that met the initial selection criteria without filtering for unique presence in a single subpopulation), valuable information about the strongest responses of several clusters to the fermentative environment emerged (Figure 3a, Figure S1, Table S4). Notably, increased expression of several genes was observed in strains from Europe/Domestic-2 and Europe-Mix (and Lt010 from Europe/Domestic-1). Highlighted genes include the two LDH genes in *L. thermotolerans* (KLTH0D00440 and KLTH0G19536), and the ortholog to *PDC5* (KLTH0D00418). These genes are crucial for central carbon metabolism, playing essential roles in converting pyruvate to lactic acid and ethanol, respectively, from pyruvate. Among them we found also a gene coding for an ABC transporter potentially involved in the transport of monocarboxylic acids, orthologous to *YBT1* (KLTH0B00396).

Conversely, we observed a distinct response involving genes related to *VEL1*. In *L. thermotolerans*, several genes are orthologs to *VEL1* in *S. cerevisiae*, among them KLTH0F00176 and KLTH0H16390. Specifically, KLTH0F00176 was upregulated in strains from the Americas subpopulation, while KLTH0H16390 showed increased expression in strains from the Europe-Mix subpopulation. This contrasting behaviour suggests that the functions encoded by these genes are differently regulated depending on the subpopulation. In *S. cerevisiae*, *VEL1* expression is induced by zinc depletion and is related to DNA replication stress and growth regulation among other processes. This suggests that *L. thermotolerans* may have adapted its gene regulation in response to environmental pressures specific to each subpopulation.

Among the unique 104 CSS, several genes related to nitrogen metabolism were downregulated in the wild subpopulations (Figure 3b, Table S4). Among them, in the Asia subpopulation, we find two amino acid permeases *MUP1* (KLTH0A03278) and *VBA2* (KLTH0E16720). The Americas subpopulation showed downregulation of a transaminase involved in ornithine degradation (*CAR2*, KLTH0E00374) and a carboxypeptidase (KLTH0H05500, *CPS1*). Meanwhile, the Canada-trees subpopulation exhibited a decrease in the expression of an oligopeptide transporter (KLTH0E01496, *OPT1*).

Additionally, the Asia subpopulation showed significant downregulation of genes involved in key steps of alcoholic fermentation, such as pyruvate decarboxylase enzymes ( KLTH0D00418, *PDC5* and KLTH0D08272, *PDC1*) and genes related to monocarboxylate permeases (KLTH0E16808, *ESBP6*). Conversely, the Europe/Domestic-2 subpopulation highlighted for the downregulation of genes involved in flocculation (KLTH0F19888, *FLO1*) and the upregulation of copper homeostasis genes through its transport across the plasma membrane (KLTH0D15004, *CTR3*). These patterns underscore the diverse adaptive strategies employed by different subpopulations in response to their environmental pressures.

### 3.4. Glucose and lactic acid genes correlation and QTTs analysis

Both the analysis of DEGs and CSS revealed significant differences in glycolysis and pyruvate metabolism, including alcoholic fermentation and lactic acid metabolism. To explore these differences further, we specifically analysed the expression levels of all genes involved in glucose catabolism in our set of 23 transcriptomes. We selected genes involved in glucose degradation (glycolysis and fermentation) as well as those involved in the subsequent mitochondrial processing of pyruvate (tricarboxylic acid cycle and electron transport chain). Lactic acid production is a specific trait of *L. thermotolerans* that may play an essential role in adaptation to the wine environment. Therefore, we performed several analyses to characterize the regulation of the genes linked to lactic acid production. Specifically, we conducted correlation analyses of the three genes encoding lactic acid enzymes (KLTH0D00440, KLTH0G15936, and KLTH0G159558) with the rest of the transcriptome and, more specifically, with those genes linked to glucose catabolism. Additionally, we performed a QTT analysis to determine the genes associated with lactic acid production by correlating lactic acid concentration in the fermentation at the time of RNA-seq sampling with the whole transcriptome, employing a previously defined and verified regression model for phenotype-gene expression linking (Kong et al., 2022; Pang et al., 2019).

LDH coding genes showed significant correlations with other genes across the general transcriptome (Figure 4a, Table S5). The strongest correlation was observed between an LDH-coding gene and a PDC-coding gene (KLTH0D00440 and KLTH0D00418, respectively), with a correlation index of 0.96. Another notable correlation was between an LDH gene and a serine/threonine phosphatase gene (KLTH0G19536 and KLTH0E03960, respectively), with the latter’s expression being induced by ethanol and osmotic stress. Other important genes positively linked with several of the LDH coding genes include those involved in nitrogen metabolism, such as transaminases similar to Aro8 and Aro9 from *S. cerevisiae* involved in amino acid synthesis and several aroma compound biogenesis (KLTH0G19624, YER152C) and vacuolar aminopeptidases (KLTH0D05544, *LAP4*), as well as those negatively correlated, such as genes involved in aromatic amino acid synthesis (KLTH0G02816, *ARO1*). Additional genes of interest include cell surface-related genes such as flocculins (KLTH0C11924, *FLO5*) and zinc-regulated surface-anchored GPI-proteins (KLTH0D00308, *ZPS1*; KLTH0H16390, *VEL1*). as well as ion and monocarboxylic acid transporters (KLTH0F04796; KLTH0G02816, *MCH2*) and pyrimidine salvage pathways (KLTH0E15290g, *FCY1*), all of which showed positive correlations.

**Figure 4.**
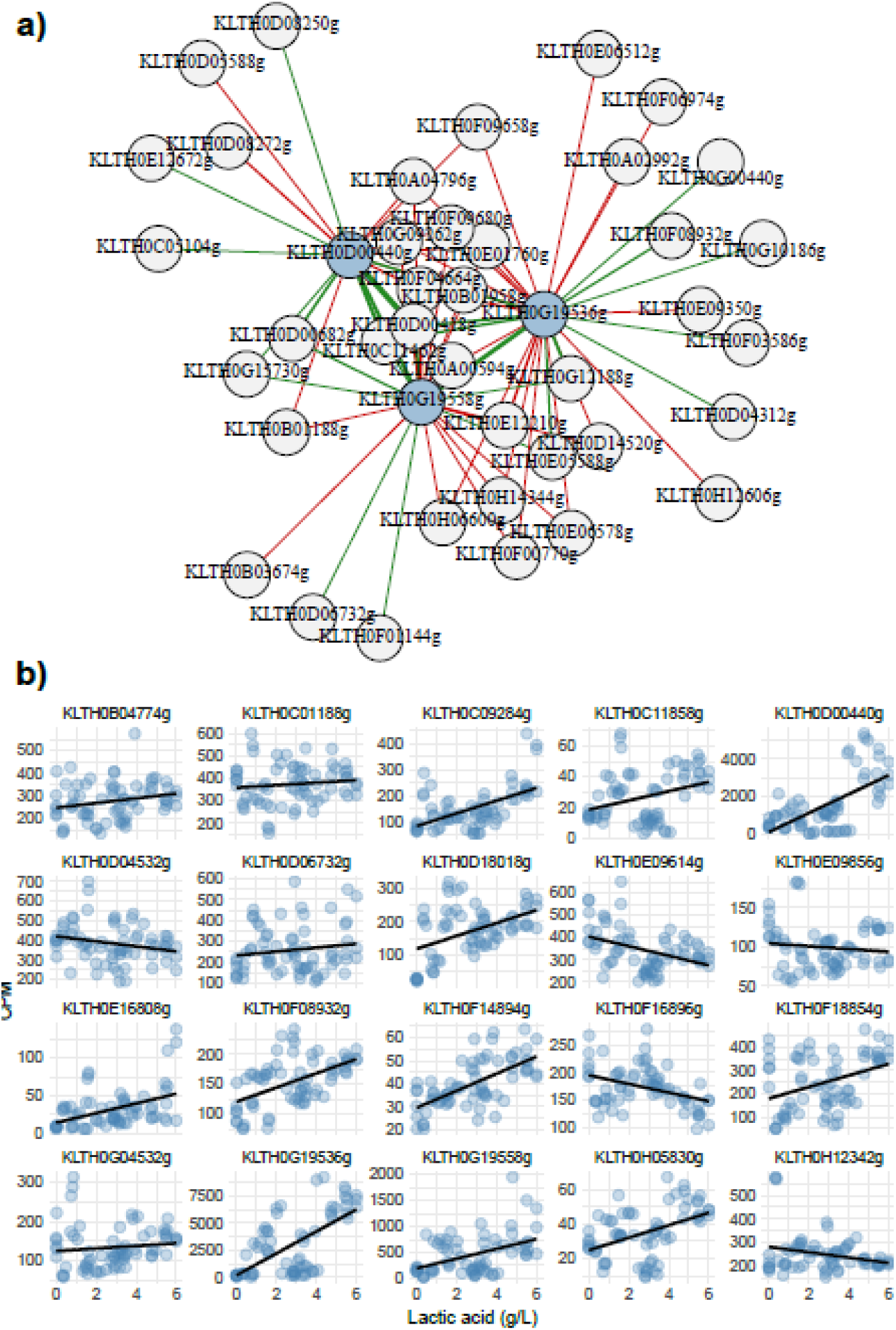
Specific relationships between lactic acid metabolism and other transcriptomic features. a) Network constructed based on the top 25 genes (grey nodes) with the highest absolute values of significant correlations (p<0.05) with the three different LDH genes (blue nodes) in *L. thermotolerans*. Edge colours represent positive (green) or negative (red) correlations, and edge width represents the correlation coefficient. b) Quantitative Trait Transcript (QTT) analysis. Counts per million (CPM) of selected genes showing significant correlations with lactic acid levels are plotted against lactic acid concentrations for all samples, including trend line.

The most relevant information, however, could be obtained by analysing the correlation of LDH coding genes with genes involved in glucose metabolism and linking phenotype to gene expression levels through QTT analysis (Figure 4b, Table S6). This analysis determines how these genes interact and whether their regulation is synergistic, as well as identifies which genes are directly involved in lactic acid production. We found a direct and negative correlation of several LDH coding genes with oxidative phosphorylation of glucose pathways (TCA and electron transport respiratory chain). Among them, genes related to the respiratory chain such as ubiquinol-cytochrome reductase (*QCR2*, KLTH0A00594; *QCR4,* KLTH0B03674; *QCR7,* KLTH0F00770; *QCR9,* KLTH0E01790; *RIF1,* KLTH0H14344) and cytochrome *c* genes (*COX9,* KLTH0D14520; *COX4,* KLTH0E06578), and genes involved in TCA such as ketoglutarate dehydrogenase (*KGD4*, KLTH0G09262) and pyruvate carboxylase (*PCY1/2,* KLTH0H06600). Conversely, specific genes involved in glycolysis and alcoholic fermentation, including lactic acid production, are positively correlated with LDH genes. These include LDHs that correlate with each other, indicating synergistic activity, and other relevant genes such as pyruvate decarboxylase (*PDC5,* KLTH0D00418) and alcohol dehydrogenase (*ADH9,* KLTH0E05588), with high correlation indices (up to 0.95). Additionally, there is a positive correlation with key glycolytic processes involving glucokinase (*GLK1,* KLTH0F01144) and glyceraldehyde 3-phosphate dehydrogenase (*TDH2/3,* KLTH0G15730), essential for initiating and continuing glycolysis, respectively. It is also notable that specific TCA enzymes, such as malic acid enzyme (*MAE1,* KLTH0D06732), fumarase (*FUM1,* KLTH0C11462), and α-ketoglutarate dehydrogenase (*KGD2,* KLTH0G12188), are positively correlated with LDH genes.

Among the genes positively correlated with the different LDH genes, several were also confirmed as QTTs of lactic acid production, demonstrating a positive correlation with lactic acid concentration at the sampling time (Figure 4b, Table S5-S6). These include genes involved in alcoholic fermentation (e.g., PDC and ADH) and the fumarase gene from TCA. Additionally, other genes acting as QTTs of lactic acid production and associated with the central carbon metabolism were identified, such as phosphoglucomutase and 6-phosphofructo-2-kinase (*PGM2* and *PFK27*, respectively). Not only central carbon metabolism-linked genes are QTTs of lactic acid production, but also genes involved in nitrogen metabolism (*PUT4, ARO8, CHO9*), zinc and iron homeostasis (*IZH1, ZPS1, ZRT1*/*COT1, FET4*), nuclear transport (*DBP5, NSP1, SAC3*), pseudohyphal growth/starvation (*STE7, PLC1*), stress response (*SSA3, RSV167, SKO*), and several transporters related to monocarboxylate acids (*AQR1, MCH2*).

### 3.5. Fermentative kinetic, metabolite production and final wines composition

The anthropization of the environment has significantly influenced the evolutionary process of *L. thermotolerans* and thereby affecting both the expression and phenotypic characteristics under winemaking conditions. To gain a comprehensive understanding of the fermentative metabolism we conducted an extensive characterization of the fermentative performance of several representative *L. thermotolerans* strains, determining not only their fermentative capacity and its impact on the chemical composition of the resultant wines (analysing major metabolites and the volatile fraction) but an examination of fermentative and population kinetics, as well as the production profiles of specific metabolites.

All tested strains were able to both proliferate and ferment properly with no significant differences observed between subpopulations with similar fermentative kinetics among the different strains (Figure S2). At the sampling time for RNAseq analysis all strains were in the exponential growth phase. Additionally, at 30 hours, all strains, regardless of their subpopulation origin, have reached between 5.2 x 10^7^ and 9.1 x 10^7^ CFU/mL, viable counts that were maintained up to the 72 hours of the fermentation.

As far as sugars are concerned, all strains followed a similar consumption kinetic. Out of the initial 180 g/L of fermentable sugars at the beginning of the fermentation, all strains had consumed between 45.7 and 52.0 g/L by the sampling time (Figure 5a). At the end of the fermentation, the mean concentration of residual sugars was 25 g/L, with specific strains leaving from 10 to near 60 g/L (Figure 5, Figure S3, Table S7). On the contrary, lactic acid production kinetic, despite following a similar pattern across all the studied strains that produce lactic acid, showed extremely different values both at 30 hours and at the end of the fermentation (Figure 5a, Figure S3-S4). Different subpopulations could be classified into three different groups regarding lactic acid production: Europe/Domestic-2 and Europe-Mix produced a final concentration around 7.5 g/L, from which, near 4.5 g/L were produced at the timepoint of RNAseq sampling (30 hours); Europe/Domestic-1, Canada-trees and Americas reached final values around 3.5 g/L (around 2 g/L at 30 hours); and Asia subpopulation reached mean final values around 1 g/L (Figure 5c, Figure S3-S4).

**Figure 5.**
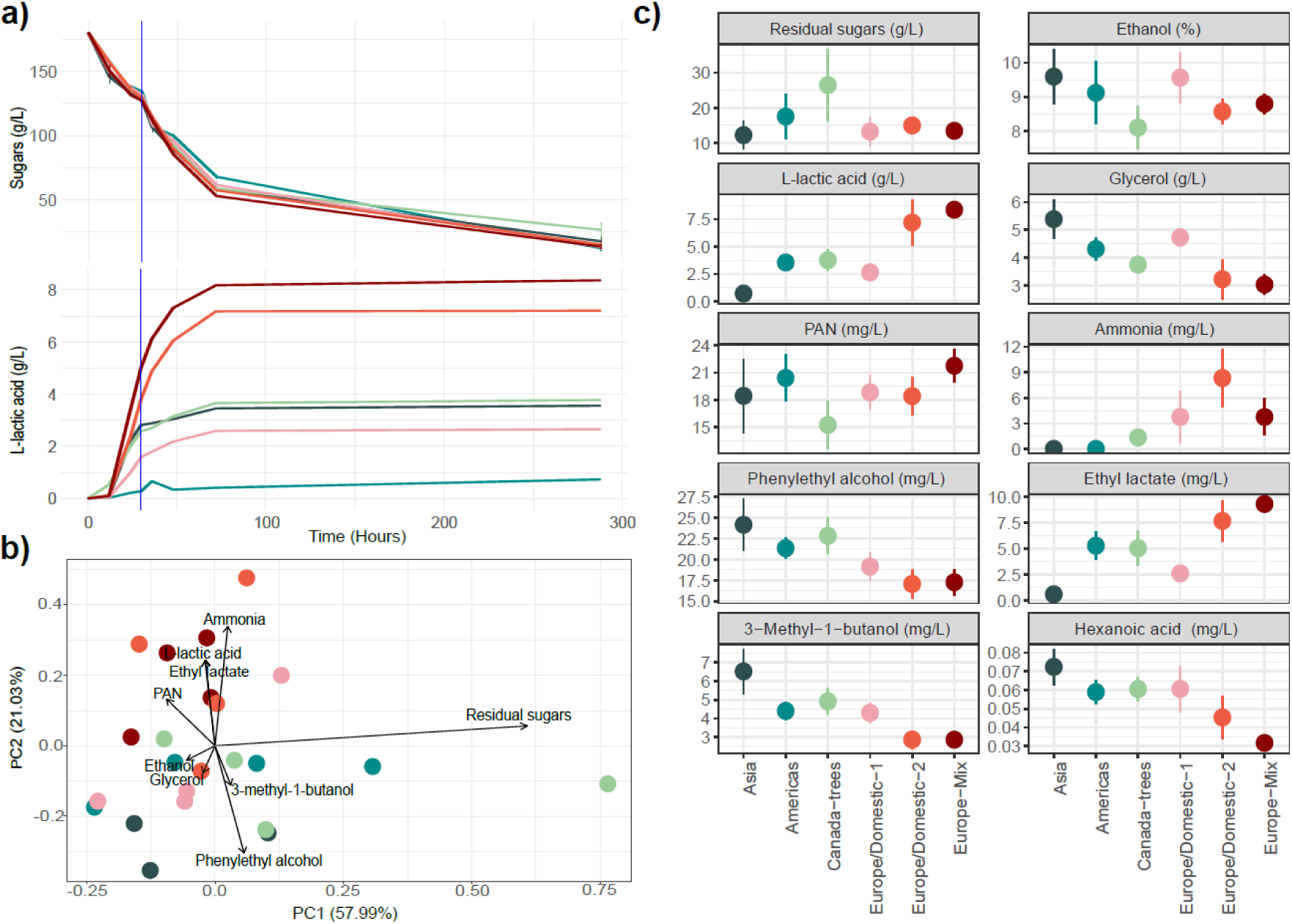
Wine chemical profile influenced by *L. thermotolerans* in laboratory-scale fermentations. a) Mean sugar consumption and lactic acid production kinetics by subpopulation. The sampling time at 30 hours is indicated by the blue vertical line. b) PCA analysis based on the final chemical profiles of wines fermented by different strains. Mean values of all metabolites measured at the end of fermentation (288 hours) for each sample were used for the calculation. The top 10 contributing variables are highlighted. c) Concentrations of selected metabolites at the end of fermentation, shown by subpopulation. The colour code for the subpopulations is consistent with Figure 1.

Differences in lactic acid production were negatively correlated with glycerol concentration at the end of the fermentation (rho= -0.85, p-value=3.84e-7), while this metabolite was positively correlated with ethanol (rho= 0.66, p-value=5.80e-4) (Figure S5, Table S8). The variation in metabolite production in the fermentation medium reflects the impact of the subpopulation origin on the resulting wine (Figure 5b-c, Figure S4). The Europe/Domestic-2 and Europe-Mix subpopulations exhibited more similar profiles at the end of the fermentation. These profiles were characterized by increased levels of L-lactic acid, which directly impacted total acidity and pH, and reduced levels of glycerol and ethanol. Other parameters did not show significant variation between subpopulations. Acetic acid, despite being linked to the pyruvate metabolism, did not show statistical differences among subpopulations. The synthetic medium used contained two different nitrogen sources: inorganic nitrogen, in the form of an ammonium salt, and organic nitrogen in the form of free amino acids. However, no statistical differences regarding its consumption were observed between subpopulations at different times for either form of nitrogen.

Regarding the volatile metabolites produced by the different yeast strains, several compounds were differentially produced among the various subpopulations (Table S9). Anthropized strains from the Europe/Domestic-2 and Europe-Mix clusters, particularly those with high lactic acid production, were characterized by a volatile profile rich in esters derived from lactic acid, such as ethyl lactate and isoamyl lactate, with concentrations reaching around 10 mg/L (Figure 5b-c, Figure S6). In contrast, these groups of strains produced lower amounts of higher alcohols, such as phenylethyl alcohol and isoamyl alcohol, with concentrations reduced by 2-to 4-fold compared to wild strains from the Asia or Americas subpopulations. A similar trend was observed in the production of fatty acids, with compounds like hexanoic acid showing a 2-fold lower concentration in the Europe-Mix subpopulation compared to the Asia subpopulation (Figure 5c, Table S9).

## 4. DISCUSSION

Lactic acid production is the most relevant phenotypic trait of *L. thermotolerans* for winemaking. Previous studies have extensively explored its application in different types of both natural and synthetic grape musts, as well as in other fermentative matrices, such as beer must (Comitini et al., 2011; Domizio et al., 2016; du Plessis et al., 2017; Hranilovic et al., 2021; Kapsopoulou et al., 2005; Vicente, Vladic, et al., 2024). Early research on yeast intraspecific diversity, from both the genetic and phenotypic point of view, described how this trait is linked to specific groups of strains (Hranilovic et al., 2017; Hranilovic et al., 2018). Other studies have examined the differential expression levels of key genes related with alcoholic fermentation and lactic acid production in low and high lactic acid-producing strains (Gatto et al., 2020; Sgouros et al., 2020), or how nitrogen metabolism and/or nutritional stress might influence this trait (Battjes et al., 2023).

In previous works, we have defined lactic acid production as a domestication trait in *L. thermotolerans* anthropization and evolutionary process through a comprehensive analysis the whole genome and phenotype of a 145-strain collection. This was done in parallel with other well-known anthropization traits in wine-related microorganisms, such as ethanol and sulphite resistance, increased assimilation of non-fermentable carbon sources, and altered nitrogen metabolism (Vicente, Friedrich, et al., 2024).

The analysis of expression profiles from a representative dataset of *L. thermotolerans*, including several strains from each of the different subpopulations, has highlighted how the anthropization process in the winemaking environment has shaped the response to fermentative conditions. Different subpopulations have shaped distinct signatures at the transcriptomic level in response to grape must fermentative conditions, primarily by modifying the regulation and expression of central carbon metabolism. The differences in gene expression levels, which impact directly on phenotype, are a direct consequence of genetic and genomic adaptations. The evolutionary processes linked to the winemaking environment are driven by its harsh and demanding conditions. In grape must, wine-related yeasts exhibit a general tendency to increase sugar and nitrogen uptake to enhance their competitiveness, which leads to rapid consumption of resources (sugar and nitrogen), accelerated ethanol production, and depletion of limiting nutrients at higher rates (Alvarez et al., 2023).

In the case of the different *L. thermotolerans* subpopulations, the overall expression profiles are distinct. The response to the fermentative environment is most drastically impacted in wild strains, showing an important increase both in the total number and magnitude of DEGs, as a direct response to stressful conditions. Additionally, we have shown that a set of specific genes tend to be differentially expressed in anthropized and wild strains. We have applied multiple approaches, from differential gene expression analysis to functional enrichment and multiple correlation analysis (gene correlation and QTT analysis), to determine several pathways linked to lactic acid production. Through this approach, we have identified several modifications in the fermentative/oxidative metabolism of glucose, organic acids, and ions homeostasis (including cell membrane and wall modifications, pseudohyphal growth, and starvation responses), and nitrogen metabolism.

Regarding carbon metabolism, several genes act as drivers of subpopulation differentiation and subpopulation-specific responses to the fermentative environment. Wild strains from the Americas, Asia, and Canada-trees subpopulations increase the expression of genes related to sugar transporters (i.e., *HXT6*), but this increased sugar uptake is not accompanied by enhanced glycolysis or accelerated sugar metabolism through the fermentative pathway as might be expected (Monnin et al., 2024). Anthropized clusters showed increased glycolytic flux by increasing the phosphorylation of sugars and key steps of glycolysis, through glucokinase and glyceraldehyde 3-phosphate dehydrogenase activities. The increased expression of the latter leads to a reduction in the production of glycerol, which negatively correlates with the fermentation product from pyruvate and lactic acid (Monnin et al., 2024). At the same time, these anthropized strains increase the expression of genes related to pyruvate degrading enzymes such as PDC, ADH, and LDH through the fermentative pathway, faster than the oxidative one (Hagman et al., 2014).

Remarkably, we found a negative correlation of lactic acid production with the expression of COX genes through QTT analysis. These genes, involved in the electron transport chain, have been previously shown to have lower expression levels in specific subpopulations of *B. bruxellensis* linked to high ethanol environments such as tequila/ethanol production (Jallet et al., 2023). Conversely, we found a positive correlation of lactic acid production with the expression of key enzymes in the TCA cycle (i.e., α-ketoglutarate dehydrogenase, fumarase, and malic acid enzyme). The TCA pathway provides reducing equivalents used by the respiratory chain to produce energy in the form of ATP under non-fermentative conditions and participates in different amino acid biosynthetic processes. Under fermentative conditions, the TCA cycle operates not as a cycle but as two separate sets of reactions: from citrate to α-ketoglutarate through the oxidative branch, and from oxaloacetate to succinyl-CoA through the reductive one (Camarasa et al., 2011). Under these conditions, α-ketoglutarate dehydrogenase enzymes play an essential role during fermentation catalysing a non-reversible step in the TCA cycle, involving the oxidation of α-ketoglutarate to succinyl-CoA through the oxidative branch of the TCA cycle (Vuoristo et al., 2016).

Alongside the lower expression levels of LDH genes and the consequent lower lactic acid production, we found a negative correlation of both the genes and lactic acid production with the expression levels of monocarboxylic acids transporter genes (*ESBP6*, *MCH2*, and *AQR1*) in several wild subpopulations These genes function not only at the cell membrane but also at the vacuolar membrane. One of the main mechanisms by which cells cope with this acidic stress is through increasing the expression levels of these organic acid transporters, which affect cell homeostasis (Tenreiro et al., 2002; Xie et al., 2024). Inside the cell, weak organic acids are present in the anionic form, requiring a transporter to leave the cell in order to maintain intracellular pH. Once outside the cell, at a pH below their pKa, organic acids are present in the undissociated (lipophilic) form, allowing them to permeate the cell membrane via simple diffusion (Mira et al., 2010).

On the contrary, we identified other transporters related to specific ions and metals positively linked, as QTTs, to lactic acid production. Homeostasis of these elements is essential for maintaining several cellular functions. Among them, zinc, whose concentration is usually maintained at high levels, plays a crucial role in protein structure by defining the morphology of different proteins and playing a structural role. We found increased expression in high lactic acid producing strains of zinc-regulated transporters (*ZRT1*) and factors that increased its expression under zinc-deficiency conditions (*IZH1*). The role of these factors is still unknown, but they may influence cell membrane permeability (participating in sterol metabolism) or act as zinc transporters in a signalling pathway (Lyons et al., 2004). Other zinc-related genes, such as *VEL1*, are positively correlated with *LDH* expression and are also associated with specific subpopulations. Despite differences in *VEL1* gene regulation among the studied subpopulations, all of them exhibit a general overexpression of this gene. Additionally, *KLTH0G00176* and *KLTH0H16390*, both orthologs of *VEL1*, are differentially expressed across different subpopulations. In *S. cerevisiae*, this gene has been shown to be induced under zinc depletion conditions and to participate in cell aggregation and the formation of structures such as velum, as observed in a sherry wine strain (Dunn et al., 2005).

These aggregation characteristics have been extensively studied in wine strains. Wine-related isolates often show several modifications in membrane and cell wall structure and cell morphology, which allow for greater survivability under winemaking conditions. These adaptations involve greater resistance to the ethanol concentration in the medium, cell aggregation, and flocculation (García-Ríos & Guillamón, 2022). We found several genes involved in these processes that are downregulated in wild subpopulations and that are positively linked to the expression of LDH genes and lactic acid levels. Flocculation capacity provides several advantages to yeast cells, allowing them to escape from difficult environmental conditions and survive nutrient starvation. Among the genetic determinants involved in the flocculation capacity of yeasts, we found genes directly related to flocculation (i.e., FLO gene family) and other genes coding for essential membrane components such as glycosylphosphatidylinositol-proteins (GPI-proteins) (Soares, 2011; Varela et al., 2020). The increased expression levels of both FLO and GPI-protein genes are related to increased cell surface hydrophobicity, inducing flocculation processes in *S. cerevisiae* (Soares, 2011). Specifically, *FLO5* (positively linked to LDHs expression) promotes cell-cell adhesion, contributing to the formation of multicellular aggregates through adhesins (Varela et al., 2020).

During wine fermentation, yeasts can utilize ammonium, amino acids, and other secondary sources such as oligopeptides and polypeptides as nitrogen sources (García-Ríos & Guillamón, 2022; Ruiz et al., 2020). Although nitrogen is among the limiting nutrients in grape must, it plays an essential role during fermentation in regulating species competition for resources. Additionally, several pathways of amino acids degradation and/or synthesis are involved in the synthesis of various aromatic compounds that impacts wine complexity and overall quality (Godillot et al., 2022). In *L. thermotolerans*, nitrogen metabolism appears to be generally downregulated in strains from wild environments, which may reduce their competitive capacities in the winemaking environment. This downregulation affects genes typically associated with nitrogen metabolism, such as oligopeptide transporters and amino acid permeases (Becerra-Rodríguez et al., 2020; Brice et al., 2018). We identified key genes in amino acid and oligopeptide assimilation as CSS in different wild subpopulations. Among them, *MUP1*, an essential part of the amino acid sensing pathway (Brice et al., 2018), and *OPT1*, a proton-coupled symporter involved in GSH, tetra- and pentapeptide assimilation as alternative nitrogen sources (Becerra-Rodríguez et al., 2020), stand out.

These differences in nitrogen metabolization may affect phenotypic diversity in *L. thermotolerans.* Previous studies have described a lower growth rate in wild strains under nitrogen limiting conditions (Vicente, Friedrich, et al., 2024). This observation could be related not only to the variying dosage of nitrogen-related genes but also to the lower expression levels of these genes, as shown in this work. Differences in nitrogen uptake may affect not only competition capacities between wild and anthropized strains in the fermentative media, but also play a crucial role in lactic acid production. Previous studies have shown that upon amino acid nitrogen starvation, *L. thermotolerans* switches on lactate production, which may be related to the presence of a transcriptional activator regulated by amino acids (Battjes et al., 2023).

Differences in the regulation of central carbon metabolism may impact secondary pathways responsible for volatile compound biosynthesis. Previous studies have demonstrated that different subpopulations and strains affect volatile compounds, particularly esters and higher alcohols, in distinct ways (Hranilovic et al., 2018). Consequently, variations in the expression levels of several genes involved in these pathways could influence the aromatic profile of wines fermented with different strains from the defined subpopulations. We observed a clear correlation between high lactic acid-producing strains and higher concentrations of esters, especially those derived from lactate, which are often considered primary contributors to wine aroma (Lasik-Kurdys et al., 2018). These esters, particularly ethyl lactate, are highly valued for their contribution to the wine bouquet, imparting fruity, buttery, and creamy aromas, and enhancing mouthfeel sensations of roundness (Sumby et al., 2010). Additionally, we noted a significant correlation between the expression levels of various LDH-coding genes and *ARO8/9*, which may be linked to the lower concentrations of phenylethyl alcohol observed in the anthropized subpopulations. *ARO8/9* encodes an aromatic aminotransferase involved in the Ehrlich pathway, crucial for releasing aromatic compounds of interest in winemaking, such as phenylethyl alcohol (Perez et al., 2022). It has been shown that in *S. cerevisiae*, deletion of this gene can enhance phenylethyl alcohol production (Romagnoli et al., 2015).

Overall, this study underscores the significant impact of anthropization on gene expression and metabolic specialization in *L. thermotolerans*. Through comprehensive transcriptomic analysis of 23 strains representing six distinct subpopulations, we have detailed how anthropized environments have driven adaptive gene expression changes, particularly enhancing fermentation-related processes such as glycolysis and pyruvate metabolism. Notably, strains from the Europe/Domestic-2 and Europe-Mix subpopulations exhibited significant deviations in their gene expression profiles compared to wild strains, highlighting how evolutionary pressures exerted by human-driven environments have modified gene expression levels.

The upregulation of lactate dehydrogenase genes and their correlation with key metabolic pathways, such as carbon assimilation, indicate a targeted adaptation for lactic acid production. This ability may enhance the fitness of wine-related strains under fermentative conditions by increasing carbon flux, which is not limited by redox cofactors. These cofactors are regenerated through lactic acid and ethanol production under fermentative conditions, as shown by the increased expression levels of these genes. Additionally, these findings provide valuable insights into the evolutionary mechanisms shaping the metabolic diversity and plasticity in gene expression regulation of *L. thermotolerans*, emphasizing the role of anthropization.

These results not only enhance our understanding of yeast adaptation but also have practical implications for optimizing fermentation processes in the winemaking industry by leveraging the unique traits of anthropized *L. thermotolerans* strains. Disentangling the differences in gene expression linked to lactic acid production could enable the rational design of fermentation processes optimized to address climate change-related challenges in modern winemaking and to better address yeast nutritional requirements.

## ACKNOWLEDGEMENTS

Funding for this research was provided by the LowpHWine companies consortia through the CDTI project LowpHWine (IDI-20210391) and the Spanish Ministry of Science and Innovation under the VinSegCalClim project (PID2020-119008RB-I00). Javier Vicente conducted this research under a fellowship from Complutense University of Madrid (CT58/21-CT59/21). We thank Ana Rosa Gutiérrez and Pilar Santamaría from the Institute of Vine and Wine Sciences (La Rioja, Spain) for their assistance in conducting the volatile profile analysis.

## AUTHOR CONTRIBUTION

Conceptualization: J.V., D.M., A.S. Investigation: J.V., S.B., A.S. Data analysis and visualization: J.V., A.S. Funding: S.B., D.M., A.S. Writing: J.V., A.S. All authors read and approved the final manuscript.

## DATA AVAILABILITY STATEMENT

The data presented in this study can be accessed on NCBI Bioproject PRJNA1145308. Sample codes and accession numbers are given in Table S1.

