## Supplementary figures for "Subpopulation-specific gene expression in *Lachancea thermotolerans* uncovers distinct metabolic adaptations to wine fermentation"

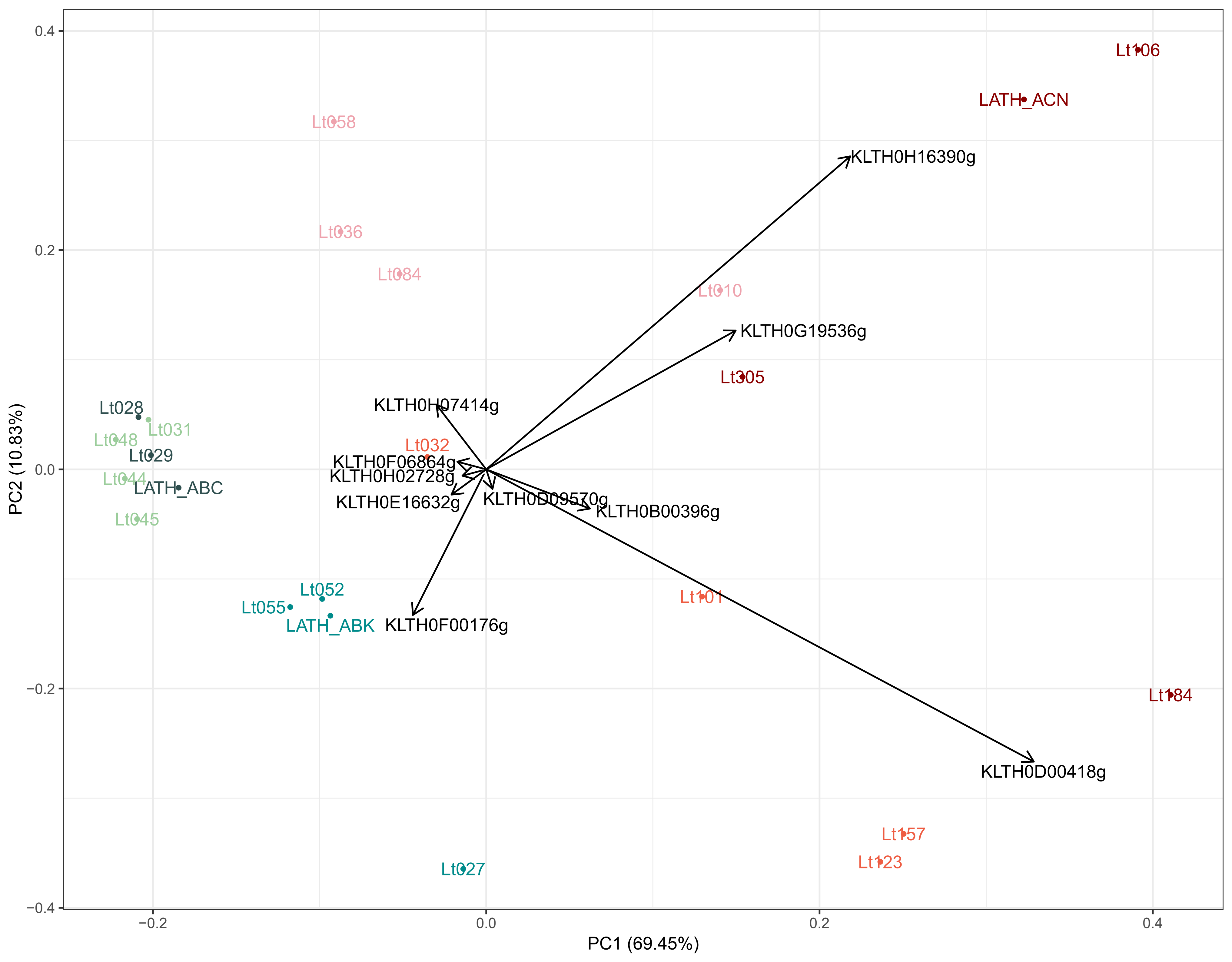

Lt106
LATH_ACN
Lt058
KLTH0H16390g
Lt036
Lt084
Lt010
KLTH0G19536g
Lt305
Lt028
KLTH0H07414g
Lt031
Lt048
Lt032
Lt029
KLTH0F06864g
KLTH0H02728g
Lt044
LATH_ABC
KLTH0E16632g
KLTH0D09570g
KLTH0B00396g
Lt045
Lt052
Lt101
Lt055
LATH_ABK
KLTH0F00176g
Lt184
KLTH0D00418g
Lt157
Lt123
Lt027
0.4
0.2
0.0
−0.2
−0.4
PC2 (10.83%)
−0.2
0.0
0.2
0.4
PC1 (69.45%)

**Figure S1.** PCA analysis of the different CSS differentiating subpopulations of *L. thermotolerans*. The top 10 contributing variables are highlighted. The colour code for the subpopulations is consistent with Figure 1.

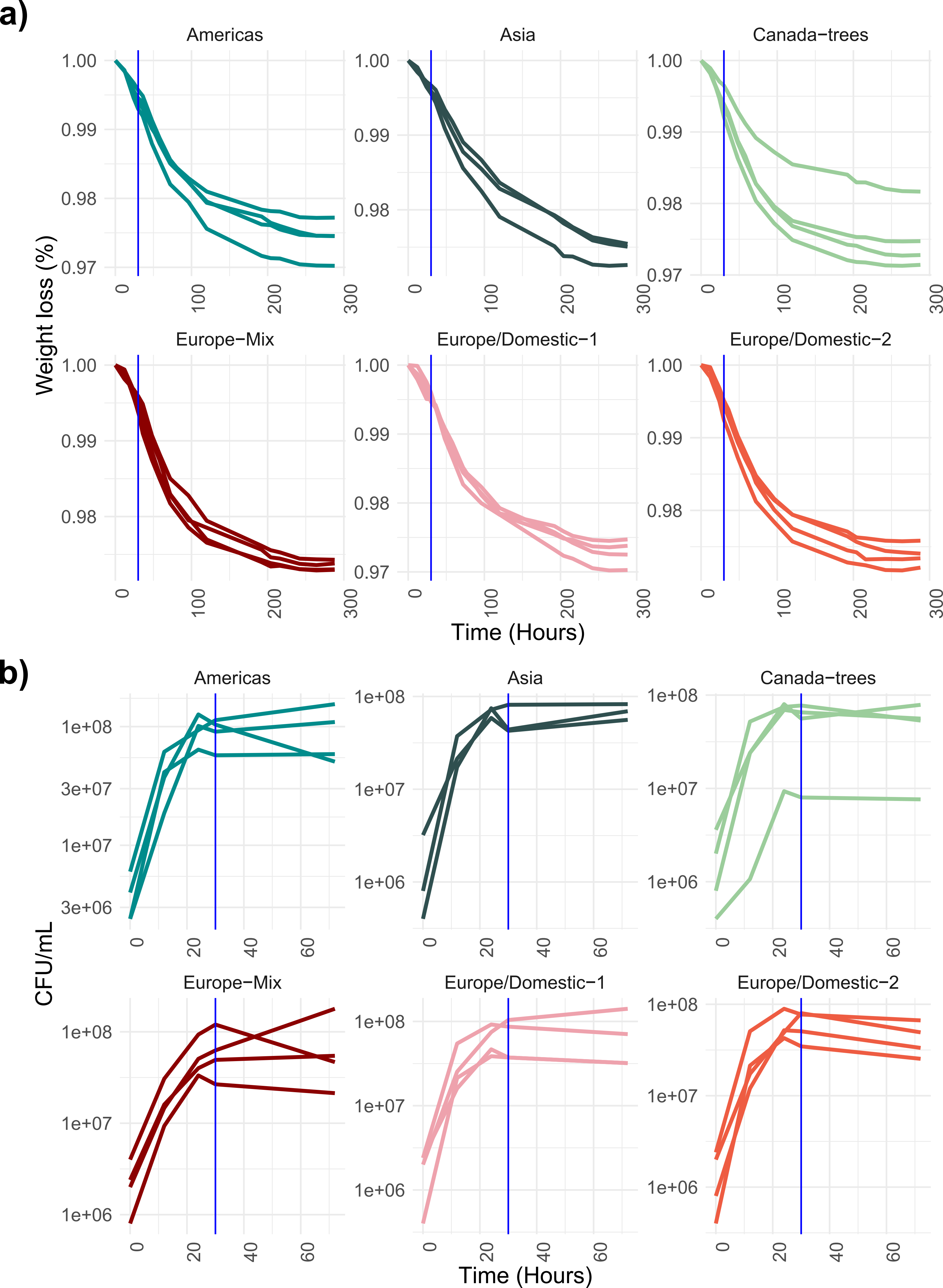
a)
Americas
Asia
Canada−trees
Europe−Mix
Europe/Domestic−1
Europe/Domestic−2
Weight loss (%)
1.00
0.99
0.98
0.97
0
100
200
300
Time (Hours)
b)
Americas
Asia
Canada−trees
Europe−Mix
Europe/Domestic−1
Europe/Domestic−2
CFU/mL
1e+08
3e+07
1e+07
3e+06
1e+06
0
20
40
60
Time (Hours)

**Figure S2.** Fermentative kinetic and population dynamics during the laboratory-scale fermentations. Weight loss (a) and viable counts (b) kinetics by each strain and subpopulation. The sampling time at 30 hours is indicated by the blue vertical line. The colour code for the subpopulations is consistent with Figure 1.

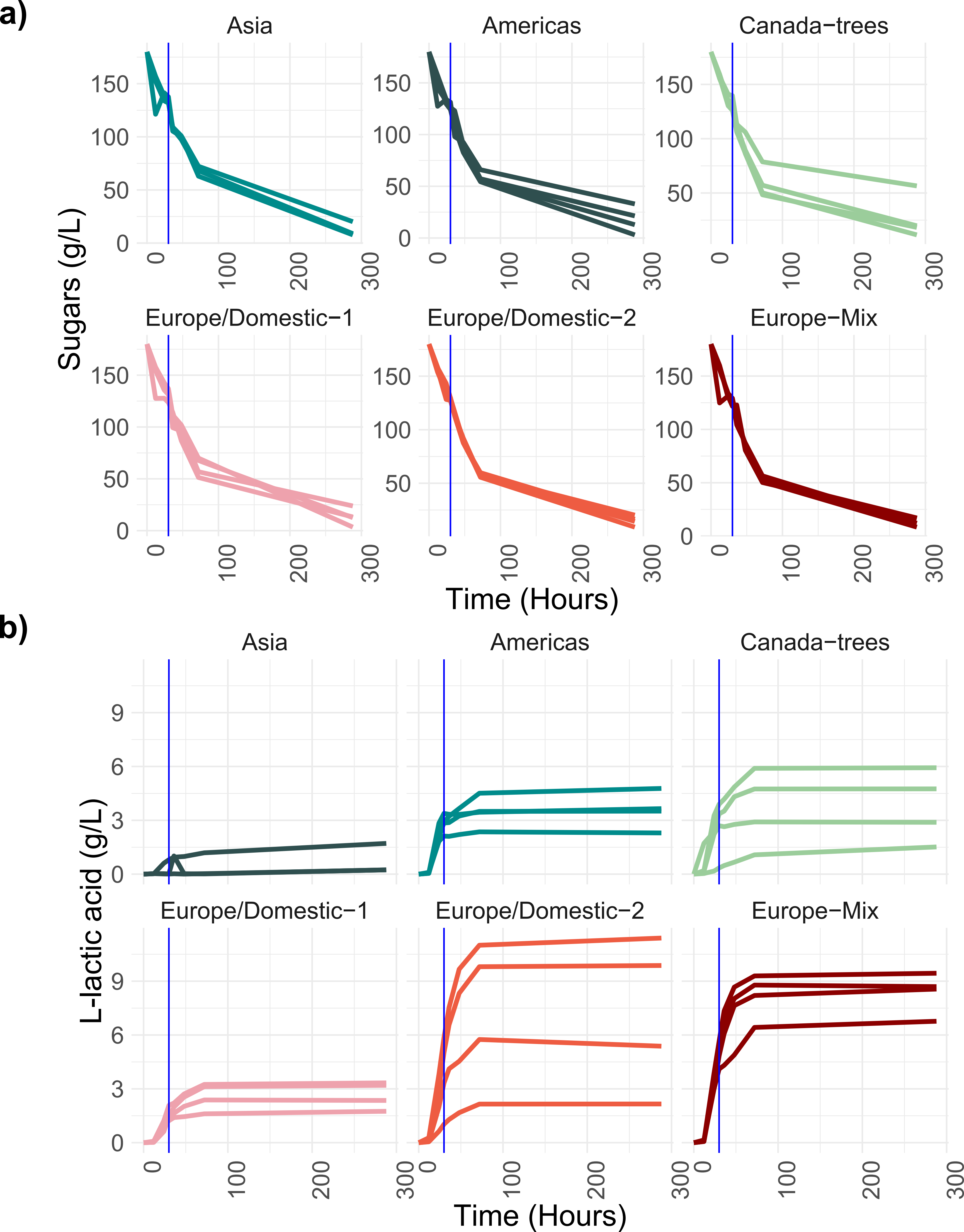

a)
Asia
Americas
Canada−trees
Europe/Domestic−1
Europe/Domestic−2
Europe−Mix
Sugars (g/L)
Time (Hours)
b)
Asia
Americas
Canada−trees
Europe/Domestic−1
Europe/Domestic−2
Europe−Mix
L-lactic acid (g/L)
Time (Hours)

**Figure S3.** Mean sugar consumption (a) and lactic acid production (b) kinetics by each strain and subpopulation. The sampling time at 30 hours is indicated by the blue vertical line. The colour code for the subpopulations is consistent with Figure 1.

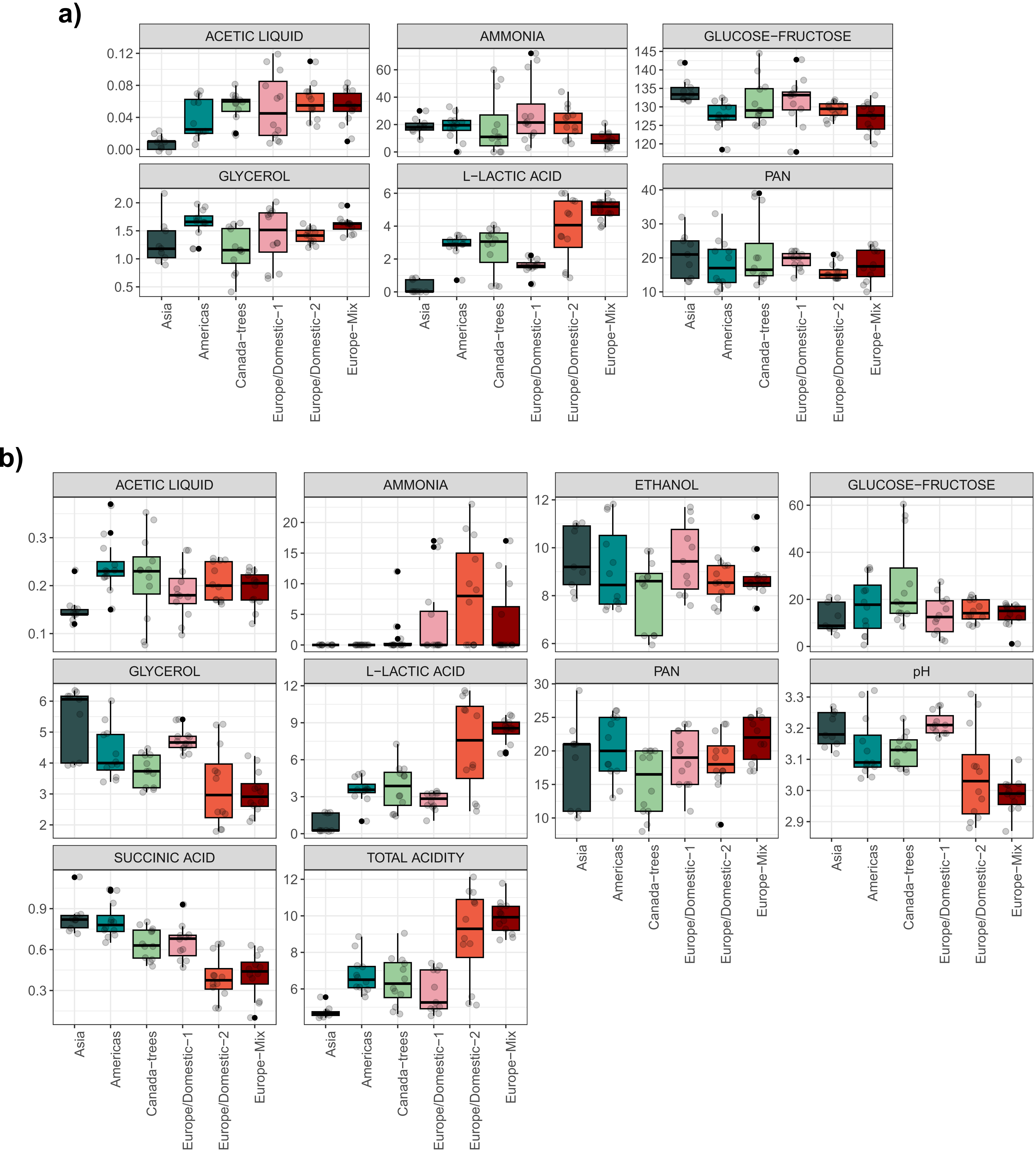

a)
ACETIC LIQUID
AMMONIA
GLUCOSE−FRUCTOSE
GLYCEROL
L−LACTIC ACID
PAN
Asia
Americas
Canada−trees
Europe/Domestic−1
Europe/Domestic−2
Europe−Mix
b)
ACETIC LIQUID
AMMONIA
ETHANOL
GLUCOSE−FRUCTOSE
GLYCEROL
L−LACTIC ACID
PAN
pH
SUCCINIC ACID
TOTAL ACIDITY

**Figure S4.** Oenological parameters measured at different timepoints by the different subpopulations and samples. a) Parameters measured at the RNA-seq sampling time (30 hours). b) Parameters measured at the end of fermentation (288 hours). All parameter units are in g/L, except for ammonia and PAN (mg/L), ethanol (%), and pH.The colour code for the subpopulations is consistent with Figure 1.

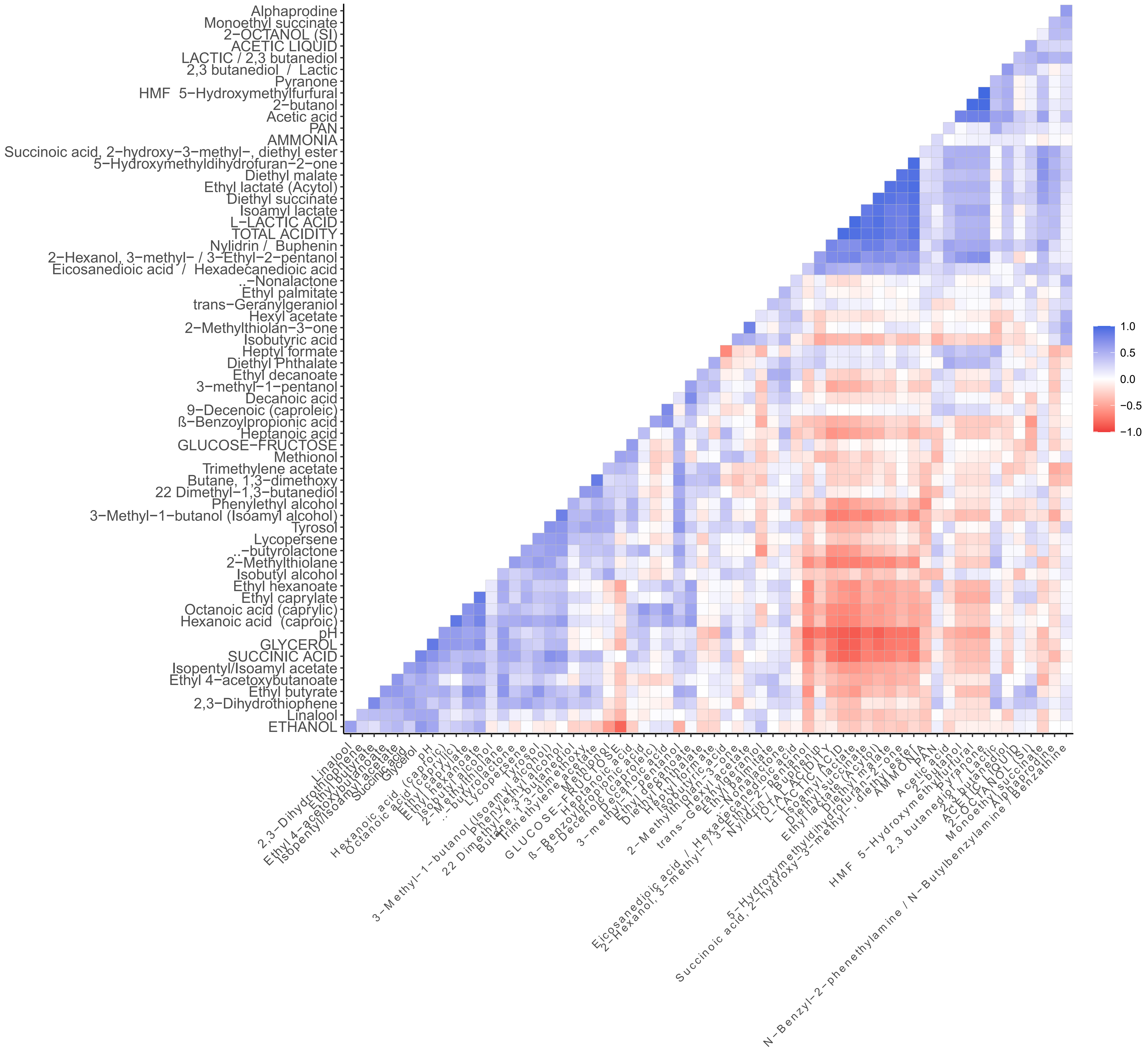

Alphaprodine
Monoethyl succinate
2−OCTANOL (SI)
ACETIC LIQUID
LACTIC / 2,3 butanediol
2,3 butanediol / Lactic
Pyranone
HMF 5−Hydroxymethylfurfural
2−butanol
Acetic acid
PAN
AMMONIA
Succinoic acid, 2−hydroxy−3−methyl−, diethyl ester
5−Hydroxymethyldihydrofuran−2−one
Diethyl malate
Ethyl lactate (Acytol)
Diethyl succinate
Isoamyl lactate
L−LACTIC ACID
TOTAL ACIDITY
Nylidrin / Buphenin
2−Hexanol, 3−methyl− / 3−Ethyl−2−pentanol
Eicosanedioic acid / Hexadecanedioic acid
..−Nonalactone
Ethyl palmitate
trans−Geranylgeraniol
Hexyl acetate
2−Methylthiolan−3−one
Isobutyric acid
Heptyl formate
Diethyl Phthalate
Ethyl decanoate
3−methyl−1−pentanol
Decanoic acid
9−Decenoic (caproleic)
ß−Benzoylpropionic acid
Heptanoic acid
GLUCOSE−FRUCTOSE
Methionol
Trimethylene acetate
Butane, 1,3−dimethoxy
22 Dimethyl−1,3−butanediol
Phenylethyl alcohol
3−Methyl−1−butanol (Isoamyl alcohol)
Tyrosol
Lycopersene
..−butyrolactone
2−Methylthiolane
Isobutyl alcohol
Ethyl hexanoate
Ethyl caprylate
Octanoic acid (caprylic)
Hexanoic acid (caproic)
pH
GLYCEROL
SUCCINIC ACID
Isopentyl/Isoamyl acetate
Ethyl 4−acetoxybutanoate
Ethyl butyrate
2,3−Dihydrothiophene
Linalool
ETHANOL
N−Benzyl−2−phenethylamine / N−Butylbenzylamine / Benzathine
1.0
0.5
0.0
−0.5
−1.0

**Figure S5.** Spearman's correlation analysis of all analysed parameters at the end of laboratory-scale fermentations. Positive correlations are shown in blue, and negative correlations are shown in red.

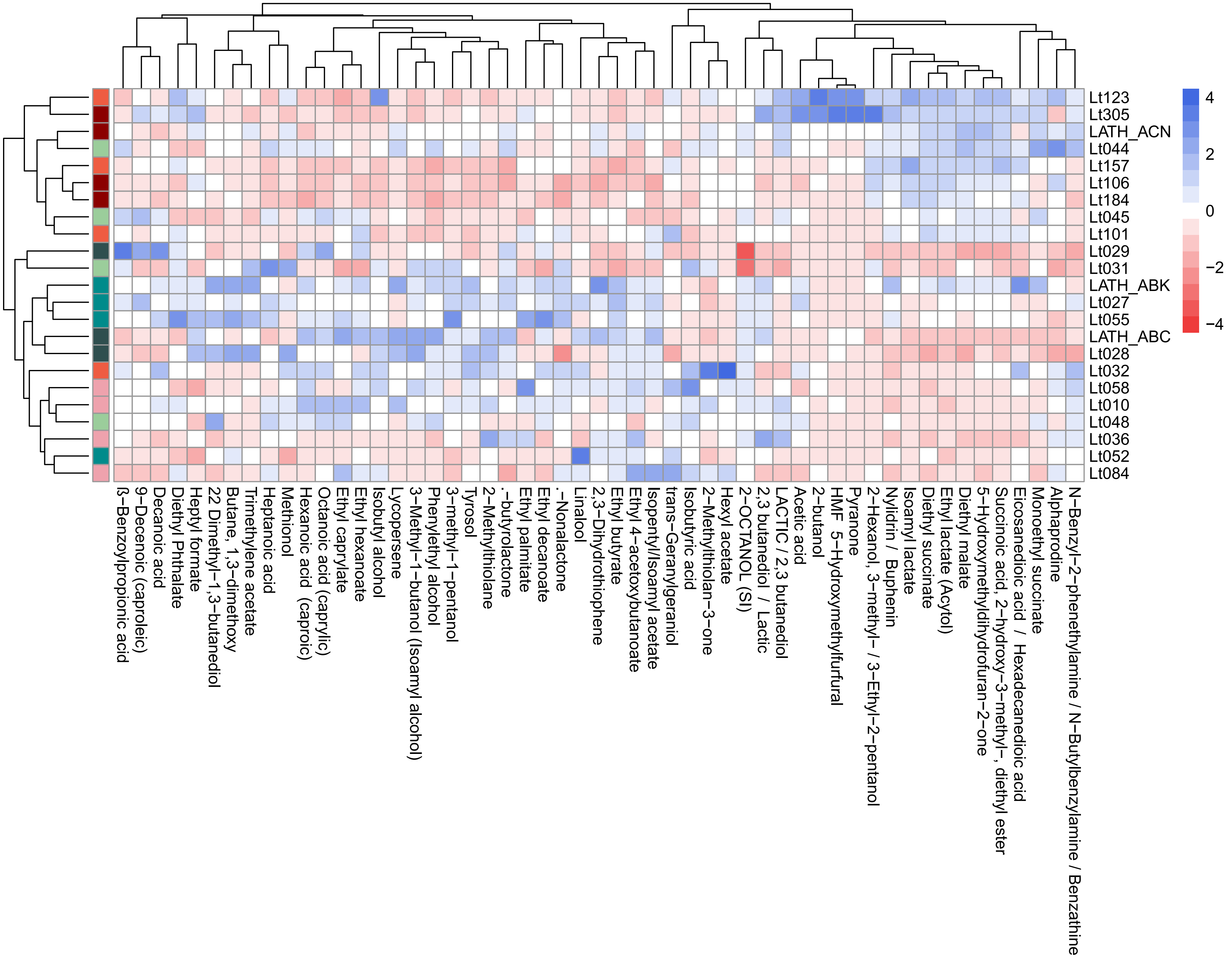

Lt123
Lt305
LATH_ACN
Lt044
Lt157
Lt106
Lt184
Lt045
Lt101
Lt029
Lt031
LATH_ABK
Lt027
Lt055
LATH_ABC
Lt028
Lt032
Lt058
Lt010
Lt048
Lt036
Lt052
Lt084
4
2
0
−2
−4
ß−Benzoylpropionic acid
9−Decenoic (caproleic)
Decanoic acid
Diethyl Phthalate
Heptyl formate
22 Dimethyl−1,3−butanediol
Butane, 1,3−dimethoxy
Trimethylene acetate
Heptanoic acid
Methionol
Hexanoic acid (caproic)
Octanoic acid (caprylic)
Ethyl caprylate
Ethyl hexanoate
Isobutyl alcohol
Lycopersene
3−Methyl−1−butanol (Isoamyl alcohol)
Phenylethyl alcohol
3−methyl−1−pentanol
Tyrosol
2−Methylthiolane
.-butyrolactone
Ethyl palmitate
Ethyl decanoate
.-Nonalactone
Linalool
2,3−Dihydrothiophene
Ethyl butyrate
Ethyl 4−acetoxybutanoate
Isopentyl/Isoamyl acetate
trans−Geranylgeraniol
Isobutyric acid
2−Methylthiolan−3−one
Hexyl acetate
2−OCTANOL (SI)
2,3 butanediol / Lactic
LACTIC / 2,3 butanediol
Acetic acid
2−butanol
HMF 5−Hydroxymethylfurfural
Pyranone
2−Hexanol, 3−methyl− / 3−Ethyl−2−pentanol
Nylidrin / Buphenin
Isoamyl lactate
Diethyl succinate
Ethyl lactate (Acytol)
Diethyl malate
5−Hydroxymethyldihydrofuran−2−one
Succinic acid, 2−hydroxy−3−methyl−, diethyl ester
Eicosanedioic acid / Hexadecanedioic acid
Monoethyl succinate
Alphaprodine
N−Benzyl−2−phenethylamine / N−Butylbenzylamine / Benzathine

**Figure S6.** Heatmap showing the differential production of volatile compounds by all strains. Mean values of each metabolite by strain were used to construct the heatmap. Data are normalized by column (by metabolite). The colour code for the subpopulations is consistent with Figure 1.

**Table S1.** List of strains employed in this work and metadata. SRAs and Bioproject accessions are included.

**Table S2.** Reads (counts) for each gene and strain determined using featureCounts.

**Table S3.** KEGG enrichment results of the DEGs by subpopulation determined using clusterprofiler. Enrichment terms were considered significant if their corrected P-adjusted value was below 0.05.

**Table S4.** CSS found in the different subpopulations. Annotation of each CSS is included.

**Table S5.** Genes showing a significant correlation with the 3 LDH genes in *L. thermotolerans*. The p-value of the correlation and the annotation of the gene is included.

**Table S6.** QTT analysis of the correlation between lactic acid production and gene expression levels. Gene codes, annotations, and KEGG functional characterizations are provided. The correlation index, p-values, and corrected p-values are shown.

**Table S7.** Main oenological parameters measured along the fermentation. Sample, analyte, time, value and units are shown.

**Table S8.** Correlation of different metabolites measured at the end of fermentation. The correlation index and p-value are shown. Correlations with p-values less than 0.05 are highlighted in bold.

**Table S9.** Volatile profile of the resulting wines fermented by each strain. Subpopulation for each strain and family for each compound are indicated. Table shows the mean ± standard deviation values.
